# The Transcriptional Gradient in Negative-Strand RNA Viruses Suggests a Common RNA Transcription Mechanism

**DOI:** 10.1101/2024.11.11.623041

**Authors:** Connor R. King, Casey-Tyler Berezin, Brian Munsky, Jean Peccoud

## Abstract

We introduce a novel model of nonsegmented negative-strand RNA virus (NNSV) transcription. Previous models have relied on polymerase behavioral differences in the highly conserved intergenic sequences. Our model hypothesizes the transcriptional gradient in NNSVs is explained through a simple model with two parameters associated with the viral polymerase. Most differences in expression can be attributed to the processivity of the polymerase while additional attenuation occurs in the presence of overlapping genes. This model reveals a correlation between polymerase processivity and genome length, which is consistent with the universal entry of polymerases through the 3’ end of the genome. Using this model, it is now possible to predict the transcriptional behavior of NNSVs from genotype alone, revolutionizing the design of novel NNSV variants for biomedical applications.

## Main Text

Nonsegmented negative-strand RNA viruses (NNSVs), including Ebola (EBOV), rabies virus, and vesicular stomatitis virus (VSV), are a diverse class of viruses that infect humans and animals, and are commonly used in biomedical applications such as vaccine vectors (*1*). Since at least 1981, it has been known that the transcription of genes in NNSVs follows a gradient where the expression of each gene is dependent on its position in the genome (*2-21*). However, the mechanism that governs the gradient has remained elusive (*20, 22*). The explanation of this gradient that has dominated in the literature is the “Start-Stop” model. This model is characterized by the entry of the polymerase at the 3’ end of the genome where it then scans down the genome until a transcriptional start signal is reached (*2-5, 23*). The polymerase then proceeds to transcribe the gene until it reaches the stop signal, at which one of two events occurs. The polymerase either continues transcribing the next gene or is ejected from the genome. The mechanism governing the ejection of the polymerase in this model has never been identified and there is no clear explanation for the variation in the ejection frequency seen at termination sequences. For example, VSV has a gradient where there is a 30% decrease in expression at each of the first 3 stop signals. However, at the 4^th^ stop signal, there is a 90% decrease in transcription. Explaining this observation requires to either assume there is a difference in the action of the intergenic regions or that intrinsic characteristics of the last gene cause this discrepancy. Currently, the commonly accepted interpretation is that the difference originates from the intergenic regions even though the sequence of the 4^th^ intergenic region is nearly identical to the other three intergenic regions.

In 2022, another model was proposed (*5*) where the authors modeled transcription as a random walk with each polymerase diffusing across the genome. The decrease in expression in the model was the result of collision-based ejections of polymerases and differences in the probability of initiating transcription at each promoter. This model proposes a few intriguing ideas that challenge some of the original assumptions of the Start-Stop model. The first is that transcription of downstream genes is not necessarily dependent on the transcription of upstream genes. In the Start-Stop model, the transcription of each gene requires the transcription of the upstream genes, which is inconsistent with real-world data as some NNSVs have overlapping genes where the transcriptional start signal of one gene is located within an upstream gene (*5, 10*). The second major idea that was introduced was that the transcriptional gradient may in part be the result of polymerases stochastically falling off the genome regardless of position.

Despite these forward-thinking ideas, both models assume that the gradient is either partly or entirely explained by differences in the action of the polymerase at stop and start sequences. However, when looking at the regulatory sequences of VSV’s genome (as well as the genes of other NNSVs), this seems unlikely as they are highly conserved with (*24*). So, it is instead more likely that there must be a factor associated with this last gene that is causing the deviation in attenuation. Additionally, the proposition of a collision-based mechanism for polymerase ejection is overly specific when we consider that it is normal for polymerases to fall off during transcription stochastically without the action of an outside force (*25-27*). In this article, we propose a novel model of NNSV transcription that challenges the traditional Start-Stop model proposed in 1981. The model suggests that the simple transcriptional gradient seen in NNSVs arises purely because of the processivity of the viral polymerase. As such, this article presents the RNA-polymerase Association Mechanism (RAM) model which proposes that most of the characteristic transcriptional gradient associated with NNSVs can be captured with a single parameter representing the probability of a viral polymerase dissociating from the genome while walking along the genome.

### Modeling Transcription

The RAM model consists of a single polymerase performing a walk beginning at the 3’ end of the genome. In each step, the polymerase takes one of two actions. The polymerase either takes a single base step forward or it falls off the genome. The probabilities associated with these actions will be referred to as p_walk_ and p_off_, respectively. Since p_off_ is the complement of p_walk_, these represent a single parameter in this model. This model has one additional parameter that is the probability that the polymerase will begin transcribing a gene when it lands on a transcriptional start signal (p_transc_). Note that the value of p_transc_ is the same for all start signals in a viral genome. The relative level of expression of most genes (p_express_) can be calculated using the equation:

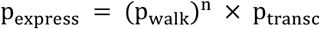

Where n is the number of base pairs that the termination sequence is from the 3’ end of the genome. It is important to note that in some NNSVs there are genes that overlap (Fig 1). In these cases, the transcriptional start signal of some genes are located within the upstream gene. In cases where overlapping occurs and the promoter of a gene (Gene 2) is located within an upstream gene (Gene 1), then the transcription of Gene 2 only occurs if the polymerase does not begin transcribing Gene 1 when the Gene 1 start signal is reached. As such, p_express_ in these cases is calculated using this formula:

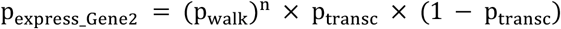

**Figure 1.**
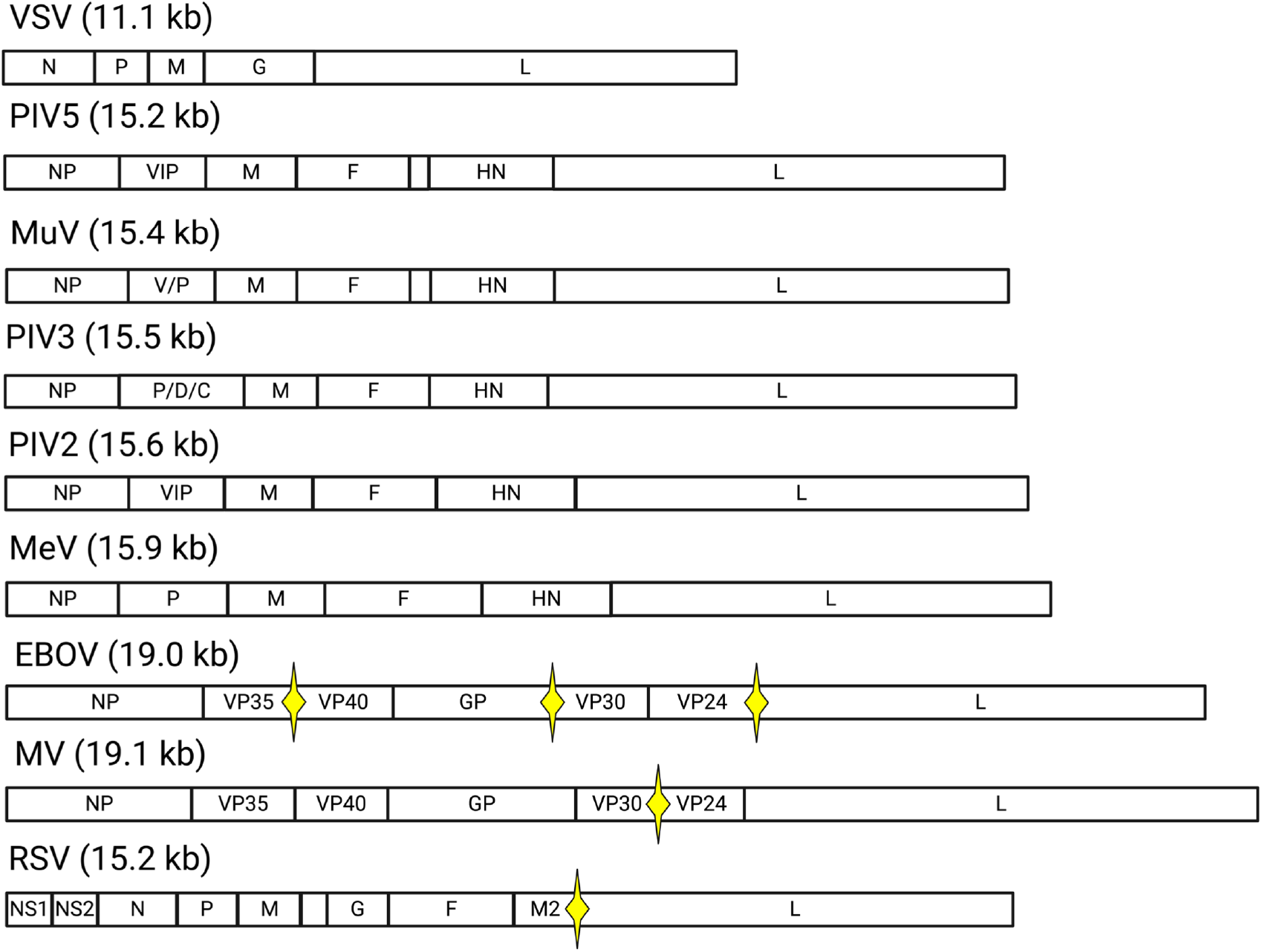
Genome Architecture of NNSVs. Graphical representation of the genomes of nonsegmented negative-strand RNA viruses (NNSVs) with gene lengths to scale. Yellow stars indicate where there are overlaps between genes in the genome. The genes that lack labels are the regions containing the small hydrophobic (SH) gene. Common proteins present in most of these species are the N/NP=nucleocapsid, P=phosphoprotein, M=matrix protein, G/GP=glycoprotein, L=RNA-dependent RNA polymerase, F=fusion protein, and HN=hemagglutinin-neuraminidase protein.

The parameters p_walk_/p_off_ and p_transc_ were determined by the fitting the model to maximize the likelihood of the data given the model, and posterior uncertainties in parameters were quantified using Markov Chain Monte Carlo (MCMC) algorithm (Table S1) (*28, 29*). The data used in this article is very diverse including sequencing data from Illumina sequencing and direct RNA sequencing as well as blotting-based methodologies. The code for fitting the models can be found in Notebook1 on the GitHub associated with this article.

### Transcriptional Attenuation at Each Intergenic Region is a Function of the Length of the Downstream Gene

The RAM model was fit to the observed gradients from 9 NNSVs of the Paramyxoviridae, Filoviridae, and Rhabdoviridae families (Fig 2, Table S2). For Rhabdoviridae, the model was fit to the original dataset used to describe the ratio of VSV expression in 1981 (*2*). For Filoviridae, EBOV and Marburgvirus (MV) were used (*10*). For Paramyxoviridae, data from Measles Virus (MeV), Mumps Virus (MuV), Parainfluenza Virus Type 2 (PIV2), Parainfluenza Virus Type 3 (PIV3), Parainfluenza Virus Type 5 (PIV5), and Respiratory Syncytial Virus (RSV) were used (*8, 30, 31*). It is important to note that the data associated with observing the transcriptional behavior of RSV are inconsistent. Some articles propose an entirely different gradient behavior of RSV with several inconsistencies when compared to the characteristic simple gradient (4, 29). In this article, the RSV gradient that most closely resembled the characteristic NNSV gradient was selected, as this is the one that is most consistent with the current understanding of NNSV transcription in literature.

**Figure 2.**
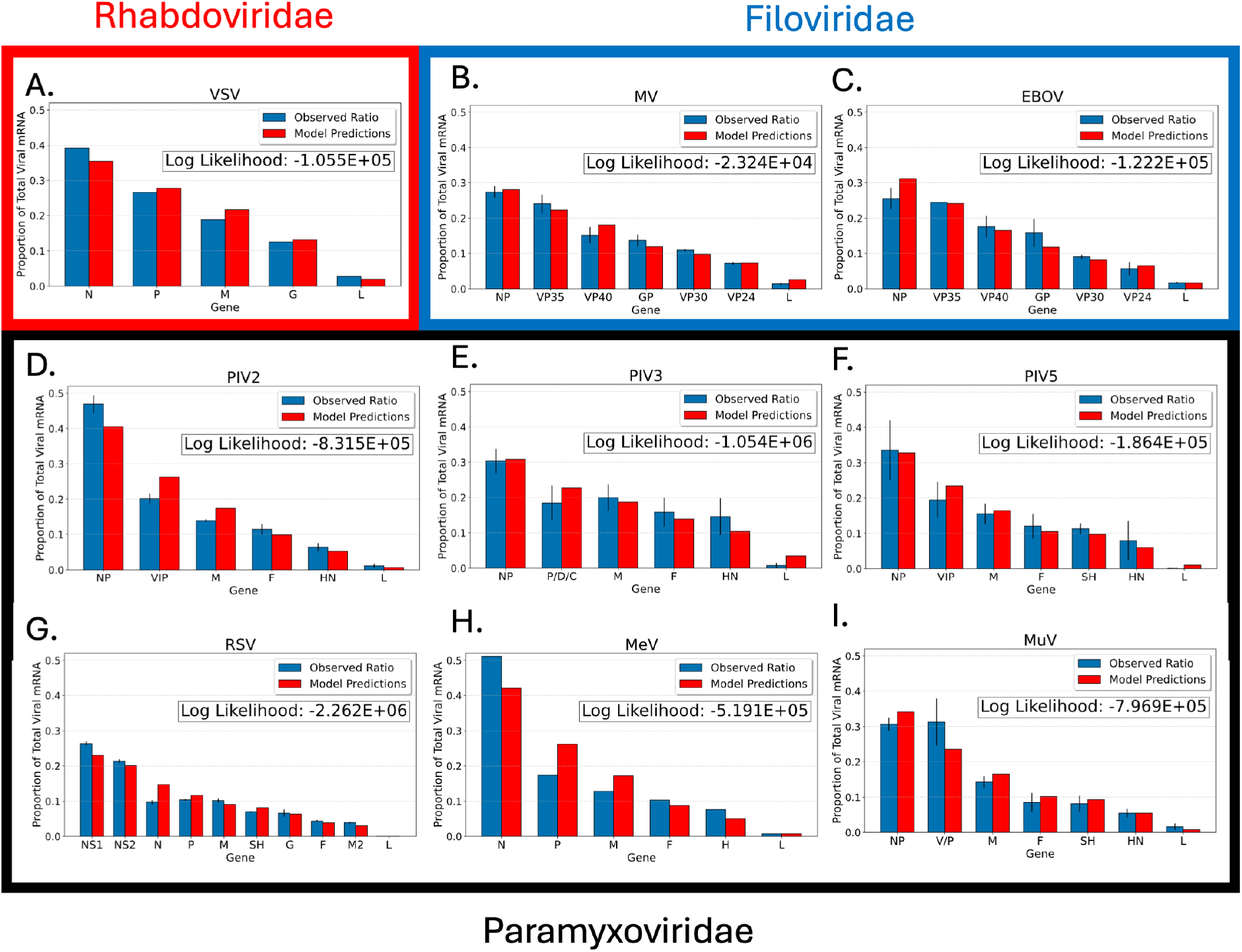
The RAM Model Accurately Predicts Transcriptional Gradients Across NNSV Species. Model predictions of the transcriptional gradient of NNSVs using the RAM model, compared to published data. These predictions are across 3 predominant families of NNSVs: Rhabdoviridae (**A**), Filoviridae (**B** and **C**), and Paramyxoviridae (**D**-**I**). Black bars are 4 standard deviations in length. When standard deviations were not available for data, there are no black bars. The log likelihood is included for every model.

The RAM model was able to achieve high-quality fits for all these viruses (Fig 2). The confidence in parameter p_walk_ associated with each virus are well-determined (Table S1). In general, the p_transc_ parameter is well-determined in the viruses that have overlaps in their genomes while it appears underdetermined in those that do not (Table S1). This is not surprising as in viruses without overlaps, all genes are equally affected by p_transc_. So, when the transcription rates are scaled to calculate the predicted gradient, this variable effectively disappears. As such, in these scenarios this model can be reduced to a single parameter. In viruses where p_transc_ is determined, it is possible to calculate the percent change in transcription rate as genes with a start sequence located within an upstream gene are attenuated further by a factor of p_transc_, since for the gene to be transcribed the upstream gene must be “missed”. There appears to be an additional 94.1%, 0.5-23.7%, and 0.3-3.4% decrease in the transcription of genes with overlaps in RSV, MV, and EBOV respectively. These results are consistent with previous findings that suggest these overlaps could be the source of inconsistencies in the characteristic transcriptional gradient of some NNSVs, especially in RSV (*5, 18, 30, 32-36*). These results suggest that the presence of overlapping genes could result in additional attenuation of transcription that is not explained by the traditional models of NNSV transcription alone. It is also important to note that this model is not inconsistent with the observation that the gradient of expression is only observed when quantifying different transcripts and not when quantifying the prevalence of sequences within the same transcript (*2*). The truncated transcripts may be rapidly degraded because they are not fully processed (*8, 37*).

### Genome Size is Correlated to Polymerase Processivity

Since p_off_ is the probability of the polymerase falling off the genome at any point, it can be used to estimate the processivity of polymerases by calculating the mean number of nucleotides added by each polymerase before falling off the genome. The highest processivity appears to be associated with MV which is predicted to travel an average of 6,599 bases along the genome, while the lowest processivity was associated with MeV which only travels about 3,331 bases on average. These values are consistent with polymerase processivities seen in other viruses (*38-40*). Intriguingly, there appears to be a negative correlation between the value of p_off_ and the length of the genome (Fig 3). This correlation suggests that for every base increase in the genome, the probability of the polymerase falling off every step down the genome decreases by about 1.79 × 10^−8^ ± 1.72 × 10^−8^ (p-value = .044, t-test statistic = 2.45). This observation is reasonable as all polymerases must enter the 3’ end of the genome and so the polymerase must stay associated with the genome for longer stretches in order for genes at the 5’ end of longer viral genomes to be expressed.

**Figure 3.**
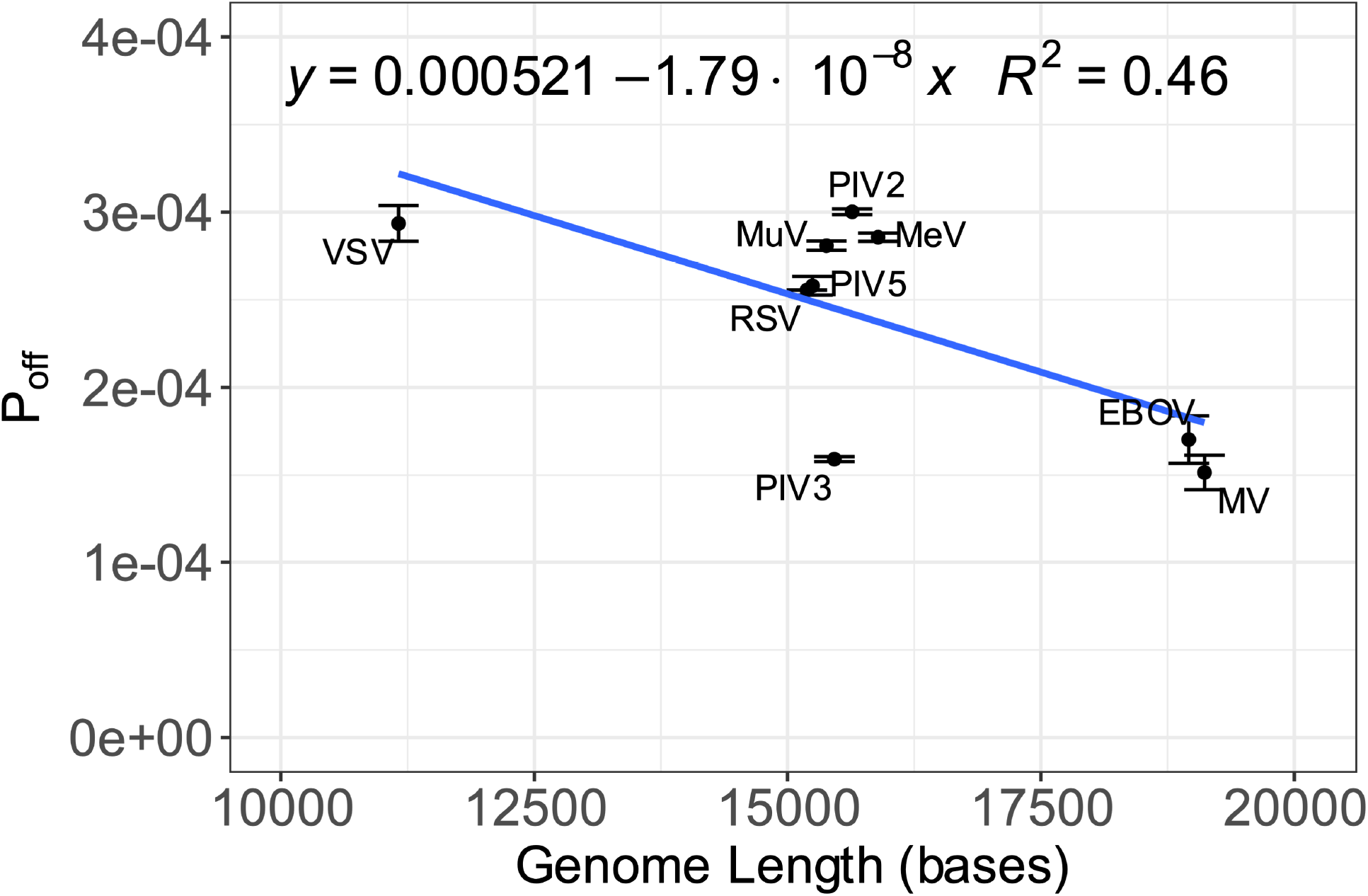
The Processivity of Viral Polymerases is Correlated to Genome Length. This figure shows a linear model showing how the probability of the polymerase falling off the genome at each step (P_off_) decreases as the length of the genome increases. The estimated slope (−1.79 × 10^−8^ ± 1.72 × 10^−8^) and intercept (5.21 × 10^−4^ ± 2.75 × 10^−4^) are both statistically significant (p-value = 0.044 and .003 respectively). The dots and error bars are the estimates and 95% confidence intervals respectively for the value of p_off_ for each virus.

### The RAM Model Accurately Predicts Alterations in Gene Expression as a Result of Genome Reorganization

Historically, VSV has been studied using “gene-shuffled” variants where the position of genes is shifted to leverage the transcriptional gradient to predictably alter transcription rates (*12, 15*). Here, the RAM Model was then evaluated on its ability to predict how shuffling the position of VSV’s genes alters the observed expression ratio of the genes (*12, 13*). The available data is all at the protein level, however, it is generally understood that the gradient in the expression of VSV genes at the transcript level also occurs at the protein level with some potential to deviate due to other mechanisms that may be controlling protein expression (*2, 12, 13, 15*). The parameters for the RAM model to generate predictions were the same used to generate Figure 2A. The RAM model was then used to predict the expression level of each gene associated with each variant from these two datasets (*12, 13*) (Fig 4A and 4B).

**Figure 4.**
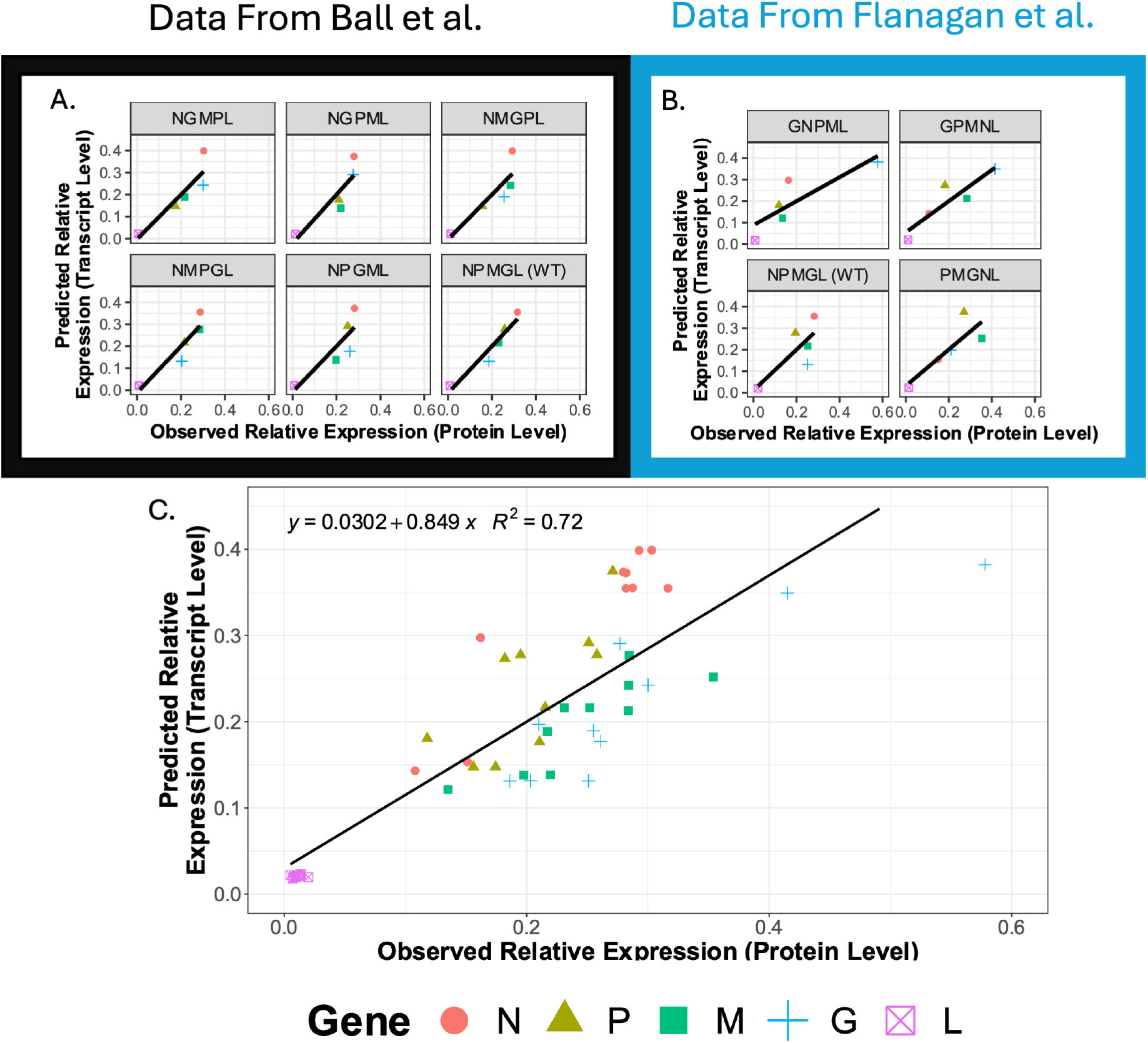
The RAM Model Accurately Predicts Transcriptional Gradients Across NNSV Variants. These plots show the correlation between the RAM model predicted expression level (based on transcription) and the observed expression level of proteins for individual gene shuffled VSV variants from 2 previous publications (A) 10.1128/jvi.73.6.4705-4712.1999 and (B) 10.1128/jvi.74.17.7895-7902.2000. (C) shows the linear fit for all variants from both datasets. The slope (0.849 ± 0.152) is statistically significant (p-value = 4.78 × 10^−15^, t-test statistic = 11.242) while the intercept is not.

It can clearly be seen that there is a positive correlation between the model predictions and the observed relative expression of genes across these variants (Fig 4C). The slope (0.849 ± 0.152) is statistically significant (p-value = 4.78 × 10^−15^, t-test statistic = 11.242) while the intercept is not. Notably, the confidence interval for the slope includes the value of 1. This is consistent with the previous statement that the expression gradient is carried on to the protein level and may occur at a nearly 1:1 level. However, it is likely that there are deviations since these proteins do not localize to the same part of the cell and so it is unlikely that they are under the exact same cellular conditions. Other environmental factors, such as the degradation rate, are likely to be different among these proteins causing some of the deviations in the data (*6*).

### Controlling Gene Order, Overlaps, and Lengths Allows for Precision Control of Viral Transcription

The model proposed in this paper proposes a fundamentally different mechanism to the traditional Start-Stop model that suggests most of the transcriptional gradient of NNSVs can be explained by their genome architecture. The evidence for this can be seen in the quality of our fits to the data across viral species (Fig 2) and NNSV variants (Fig 4), the consistency of our model with the highly conserved nature of transcription regulatory sequences in NNSVs, and the correlation of the polymerase’s processivity with genome size (Fig 3). In many cases, the model for predicting the transcriptional gradient can be reduced to a single parameter when there are no overlapping genes present in the genome. This single-parameter model is conceptually similar to a model that was briefly mentioned previously (*8*), but is mathematically distinct as the decrease in expression in the previous model was linear while the decrease in expression in this model is exponential. It should be noted that the datasets used in this article are from a diverse set of viruses and an equally diverse set of methods for characterizing these patterns ranging from traditional methods that use ^3^H-uridine to quantify viral RNAs (*2*) to next-generation sequencing data (*10*). The ubiquity of this model’s ability to predict transcriptional behaviors across methodologies and families of NNSVs and NNSV variants provides overwhelming evidence in favor of this model.

The ability to predict transcriptional behaviors based solely on genotypic information has major implications for the design of NNSV variants to both further illuminate fundamental mechanisms and meet biomedical needs. Consider a direct application towards a biomedically relevant NNSV, VSV. It has been a major point of focus to understand how the alteration of transcription rates influences the fitness of the VSV, primarily to attenuate it for medical applications (*6, 15, 41*). Considering what this article reveals about the fundamental mechanisms controlling NNSV transcription rates, there are now two additional types of VSV transcriptional variants, in addition to the historically studied gene-shuffled variants, that can be designed: gene-overlapped variants and gene-length variants. Together, these add an additional level of complexity to the control researchers can exert over transcription when designing variants. When considering gene-overlapped variants, if the presence/absence of a gene overlap is treated as a simple binary variable, this expands the total number of possible VSV variants with predictable phenotypic alterations to 1,920 possible variants when combining gene overlaps with gene shuffling.

Beyond this first simple set of variants, the possibilities grow exponentially when gene-length variants are also considered. This model suggests that the length of genes also influences transcription rates. VSV can accommodate multiple kilobases of extra bases in its genome (*42-48*). While this has historically been used to incorporate exogenous genes, this capacity could be used to further refine transcription rates by adjusting the length of genes. Genes could be potentially shortened to increase their expression if desired, as many of the genes on VSV contain additional sequences beyond the coding sequence. For example, the expression of the N gene is commonly identified as a major promoter of viral replication (*15, 41, 49, 50*). The highest possible rate of transcription for the N-encoding gene has for many years had an upper limit dictated by the transcription rate it can achieve when in the first position of the genome. The N gene contains an additional 13 bases upstream of the CDS and 44 bases downstream of the translational stop signal. If it is possible to trim this sequence, it could be possible to increase the transcription rate of the N-encoding gene and therefore increase the replicative fitness of the virus. With the proper implementation of the combination of gene-shuffling, overlapping genes, and careful design of gene lengths, the potential design space for VSV is limitless. Never has the control of transcription for VSV and other NNSVs been so customizable. Instead of relying on the natural gradient, it may now be possible to precisely alter transcription rates to obtain a refined phenotype.

Additionally, transcriptional gradients that result from the distance of a gene’s termination sequence from the entry point are not unique to viruses. Particularly, expression gradients are commonplace in long operons of prokaryotic organisms (*51*). However, these show an inverted relationship where the farther a gene’s end is from the start signal the more it is expressed at the protein level, likely due to the coupling of transcription and translation in prokaryotes. It is possible that the RAM model could be further modified to investigate these systems.

## Author contributions

Conceptualization: CRK, JP

Methodology: CRK

Investigation: CRK

Visualization: CRK

Funding acquisition: CRK, JP

Writing – original draft: CRK

Writing – review & editing: CRK, CTB, BM, JP

## Data and materials availability

All code associated with this study can be found at https://github.com/Peccoud-Lab/NNSV_Transcription_Model_2024.

