## Supplementary data for "The Transcriptional Gradient in Negative-Strand RNA Viruses Suggests a Common RNA Transcription Mechanism"

**The PDF file includes:**

Materials and Methods
Tables S1 to S2

### Materials and Methods

Code Availability

The code for this article can be found at https://github.com/Peccoud-Lab/NNSV_Transcription_Model_2024.

Data

The data used to fit the models can be found in Notebook1 and Notebook2 on GitHub. The expression data utilized comes from a variety of NNSVs including members of the Rhabdoviridae *(2)*, Filoviridae *(10)*, and Paramyxoviridae *(8, 29, 30)*. The data for sequence positions can be found in the same notebooks. The GenBank Accession Numbers of the viral genomic sequences used can be found in Table S2.

Parameter estimation

To estimate parameters, the log likelihood for observing the data based on the model-predicted ratios was maximized. The minimize function from SciPy was utilized. Each model was fit 10,000 times starting at a value between 0 and 1. The parameters chosen for downstream analyses were the ones associated with the largest log likelihood. Confidence in parameters was assessed using MCMC algorithm *(27, 28)*. The code for fitting models and performing MCMC can be found in Notebook1 on GitHub. The code for then making predictions for gene shuffled variants can be found in Notebook2.

Linear Regression

Linear regression was performed in R using the base lm function. The performance package was used to test model assumptions of normality, homogeneity of variance, and linearity. No violations were observed. The code for fitting the linear regression model connecting polymerase processivity and genome length can be found in the Markdown1 file on GitHub. The code for fitting the linear regression model connecting RAM model predicted expression and observed expression for gene shuffled variants can be found in the Markdown2 file on GitHub.

**Table S1.** Best Fit Parameter Along with Confidence Statistics from MCMC for RAM Model

| Virus | Parameter | Best Fit Estimate | Best Fit Log Likelihood | 2.5% Quantile | 97.5% Quantile |
|---|---|---|---|---|---|
| VSV | p(walk) | 0.99970 | -1.05E+05 | 0.99969 | 0.99972 |
| VSV | ptransc | 0.48017 | -1.05E+05 | 0.01917 | 0.55355 |
| MeV | p(walk) | 0.99971 | -5.19E+05 | 0.99971 | 0.99972 |
| MeV | ptransc | 0.78578 | -5.19E+05 | 0.43774 | 0.93949 |
| PIV2 | p(walk) | 0.99970 | -8.32E+05 | 0.99970 | 0.99970 |
| PIV2 | ptransc | 0.31995 | -8.32E+05 | 0.08466 | 0.38612 |
| PIV3 | p(walk) | 0.99984 | -1.05E+06 | 0.99984 | 0.99984 |
| PIV3 | ptransc | 0.07306 | -1.05E+06 | 0.07306 | 0.33737 |
| PIV5 | p(walk) | 0.99974 | -1.86E+05 | 0.99974 | 0.99975 |
| PIV5 | ptransc | 0.22064 | -1.86E+05 | 0.01927 | 0.35834 |
| MuV | p(walk) | 0.99972 | -7.97E+05 | 0.99972 | 0.99972 |
| MuV | ptransc | 0.51869 | -7.97E+05 | 0.19207 | 0.66899 |
| EBOV | p(walk) | 0.99982 | -1.22E+05 | 0.99982 | 0.99984 |
| EBOV | ptransc | 0.03440 | -1.22E+05 | 0.00314 | 0.03440 |
| MV | p(walk) | 0.99985 | -2.32E+04 | 0.99983 | 0.99986 |
| MV | ptransc | 0.10189 | -2.32E+04 | 0.00459 | 0.23717 |
| RSV | p(walk) | 0.99974 | -2.26E+06 | 0.99974 | 0.99974 |
| RSV | ptransc | 0.94093 | -2.26E+06 | 0.94093 | 0.94093 |

**Table S2.** GenBank Accession Numbers for Nonsegmented Negative-Stranded RNA Virus Genomes.

| **Virus** | **GenBank Accession** |
|---|---|
| Vesicular Stomatitis Virus | J02428 |
| Measles Virus | NC_001498 |
| Mumps Virus | JN012242 |
| Parainfluenza Virus 2 | KM190939 |
| Parainfluenza Virus 3 | NC_075446 |
| Parainfluenza Virus 5 | NC_006430 |
| Ebola Virus | NC_002549 |
| Marburgvirus | DQ447653 |
| Respiratory Syncytial Virus | M74568 |
